# High-dimensional spatial proteomics and novel machine learning pipeline identifies disease specific renal damage states

**DOI:** 10.64898/2026.01.14.699505

**Authors:** Thao Cao, Madeleine S. Torcasso, Junting Ai, Satoshi Hara, Michael S. Andrade, Anthony Chang, Gabriel Casella, Anita S. Chong, Maryellen L. Giger, Marcus R. Clark

**Affiliations:** Department of Medicine, Section of Rheumatology and Gwen Knapp Center for Lupus and Immunology Research, University of Chicago, Chicago, IL, USA; Pritzker School of Molecular Engineering, University of Chicago, Chicago, IL, USA; Department of Radiology and Committee on Medical Physics, University of Chicago, Chicago, IL, USA; Department of Surgery, University of Chicago, Chicago, IL, USA; Department of Pathology, University of Chicago, Chicago, IL, USA; Department of Medicine, Section of Hematology/Oncology, University of Chicago, Chicago, IL, USA; Department of Nephrology and Rheumatology, Kanazawa University Hospital, Kanazawa, Ishikawa, Japan

## Abstract

Lupus nephritis (LuN) and renal allograft rejection (RAR) manifest inflammation and fibrosis that ultimately lead to kidney failure. To quantitatively assess spatial injury patterns, we collected high dimensional spatial proteomics data from 23 LuN, 33 RAR, and 8 kidney control (KC) biopsies. We developed a computational pipeline to segment and classify tubules, capillaries, and glomeruli in whole-slide images using three trained neural networks (Renal Damage diagnosis, RDDx). RDDx achieved high accuracy and generalizability, reliably identifying small capillaries and differentiating tubular and vascular inflammation in kidney tissues. Both LuN and RAR showed reduced tubular and capillary areas with expanded interstitial space. LuN displayed patchy clusters of stressed and inflamed tubules, whereas RAR exhibited diffuse injury. Within RAR, T cell–mediated rejection (TCMR) showed intense tubulitis while antibody-mediated rejection (ABMR) featured proliferating and inflamed capillaries near atrophic tubules. RDDx quantitative metric outputs correlated with histopathological scores, highlighting their reproducibility and clinical relevance. Stressed tubules in mildly inflamed LuN biopsies suggested they were a sensitive injury marker, while proliferating capillaries revealed microvascular remodeling in ABMR. These findings indicated RDDx can identify and quantify damage mechanisms specific to each renal disease thus facilitating future mechanistic studies and therapeutic target discovery.

## Introduction

Tools to identify and quantify different types of renal injury remain critically needed to advance both the understanding and treatment of human kidney diseases (1,2). As examples, lupus nephritis (LuN) and renal allograft rejection (RAR) are immune-mediated diseases that damage the kidney and can ultimately lead to kidney failure or graft loss respectively (3–7). LuN is a severe complication of systemic lupus erythematosus (SLE) with multifactorial hallmark features including deposition of immune complexes, glomerular injury, tubulointerstitial inflammation, and fibrosis (4, 5). Whereas LuN is driven by autoimmunity, RAR is a complex orchestrated immune response against the kidney allograft, resulting in graft inflammation. RAR is classified into three subsets: antibody-mediated rejection (ABMR), T-cell mediated rejection (TCMR), and mixed rejection (MR) with features of both ABMR and TCMR (6). Characterizing tissue injury in these renal diseases would enable a better understanding of the relationships between different inflammatory and renal damage states and provide valuable insights for clinical assessments (8–12).

The current gold standard for diagnosis is kidney biopsy followed by microscopic examination to score inflammation, structural damage, and immune cell infiltration (11, 12). However, because histopathology scoring is performed by humans, issues arise due to subjectivity, inter-observer variability, and reliance on semi-quantitative scoring guidelines (13, 14). In addition, most criteria for staging LuN severity emphasize the pattern and extent of glomerular lesions (11) and overlook damage in other kidney structures—tubules and microvasculature—that strongly influence LuN prognosis (15, 16). Thus, in LuN and other renal diseases, comprehensive, objective, and quantitative assessments of renal injury are needed (17–20).

Recent advances in both tissue imaging and machine learning-driven data analysis have provided powerful tools to address the above research and clinical needs (21–27). Advanced imaging platforms, such as multiplexed ion beam imaging (MIBI), tissue-based cyclic immunofluorescence (t-CyCIF), and co-detection-by-indexing (CODEX), enable simultaneous capture of multiple molecular markers with spatial mapping at subcellular resolution (28–31). Machine learning architectures (e.g. U-Net and YOLO) and tools (e.g. Cellpose, Mesmer, and BANKSY) can be used to segment, classify, and phenotype hundreds of thousands of cells across large tissue areas (32–37). Collectively, these advances allow us to systematically characterize tissue architecture and injury at single-cell resolution and link microanatomical damage with complex immune landscapes (45–49).

In this study, we used CODEX multiplexed fluorescence imaging and trained three parallel convolutional neural networks to characterize inflamed and damaged parenchymal structures in LuN and RAR. We refer to the computational pipeline as Renal Damage Diagnosis (RDDx). Phenotypic classification of glomeruli, tubules, and capillaries at single-structure resolution revealed both common damage mechanisms and those unique to each disease. This quantitative framework will enable future studies to precisely identify how specific immune mechanisms are damaging the kidney in different renal diseases. Furthermore, RDDx offers a scalable pipeline for histopathological phenotyping that could be used to improve the diagnosis and staging of LuN, RAR subsets, and other renal diseases.

## Results

### A high-dimensional immunofluorescence renal dataset

To characterize kidney structures (i.e. glomeruli, tubules, capillaries) and examine their *in situ* spatial distributions and relationships, we selected and imaged 64 formalin-fixed paraffin-embedded (FFPE) biopsies using CODEX fluorescence microscopy (Fig. 1A, Table 1). The dataset consists of core-needle kidney biopsies from 23 patients with LuN and 33 patients with RAR, as well as nephrectomy biopsies from eight patients with renal cell carcinoma which served as a kidney control (KC) cohort in this study (Table 2). Among the RAR samples, nine came from patients with ABMR, nine from TCMR, and 15 from MR (Table 2). Pathological scores reported by a renal pathologist revealed no significant differences among different pathologies for chronicity index (CI), interstitial fibrosis (IF), and tubule atrophy (TA) but significantly lower tubulointerstitial inflammation (TI) scores for ABMR compared to the other diseases (Table 3). This is consistent with ABMR being associated with microvascular, not tubulointerstitial, inflammation (38, 50).

**Figure 1.**
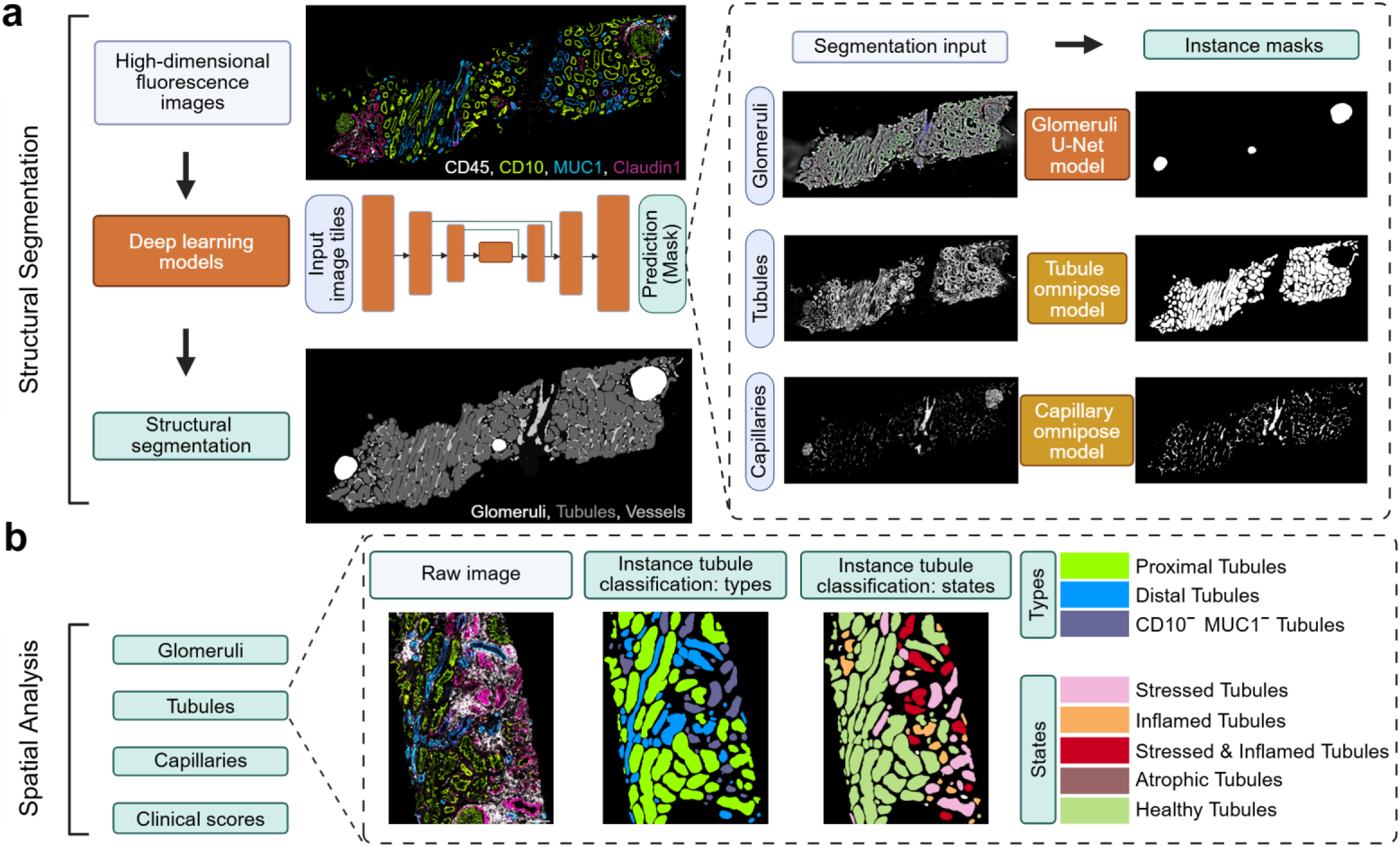
Renal structural segmentation and analysis (RDDx) pipeline. (a) High-dimensional fluorescence images were obtained and preprocessed to become input for three deep learning neural networks. These neural networks are U-Net based and trained to segment single instances of kidney glomeruli, tubules, and capillaries. (b) Combining the outputs of the networks and fluorescence biomarkers, each kidney tubule was classified by type (proximal CD10^+^, distal MUC1^+^, or double negative CD10^-^ MUC1^-^) and state (stressed Claudin1^+^, inflamed CD45^+^, stressed & inflamed Claudin1^+^ CD45^+^, atrophic CD10^-^ MUC1^-^ with high circularity, or healthy). Similarly, each capillary was classified as inflamed (vasculitis) CD45^+^, proliferating Ki67^+^, inflamed and proliferating CD45^+^ Ki67^+^, or healthy.

**Table 1.** CODEX primary antibodies and staining agent used in this study.

| Target | Clone | Manufacturer | Catalog No. | Dilution |
| --- | --- | --- | --- | --- |
| CD10 | EPR22867 | abcam | ab256494 | 1:200 |
| CD31 | EP3095 | abcam | ab226157 | 1:200 |
| CD34 | QBEND/10 | Invitrogen | MA1-10202 | 1:200 |
| CD45 | EP322Y | abcam | ab214437 | 1:200 |
| CD138 | EPR6454 | abcam | ab226108 | 1:200 |
| Claudin1 | EPR121871 | abcam | ab238949 | 1:200 |
| MUC1 | 955 | Novus Biologicals | NBP2-44658 | 1:200 |
| Ki67 | B56 | BD | BDB556003 | 1:200 |
| Nuclear staining | N/A | Invitrogen | H3570 | 1:1000 |

**Table 2.**
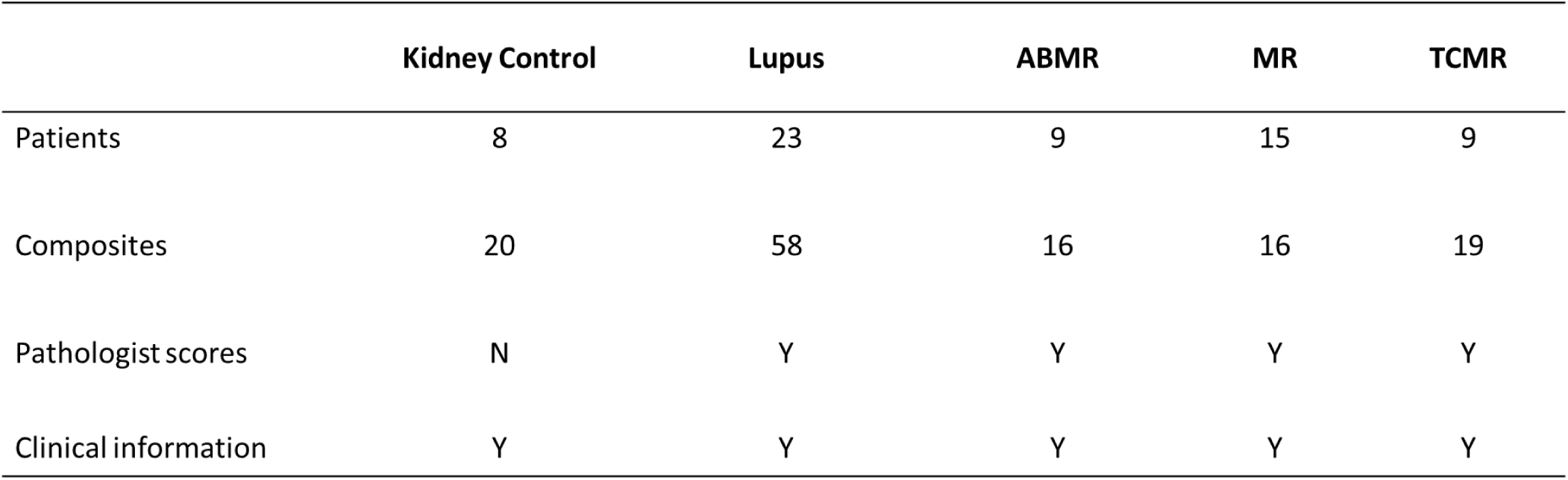
Summary of this study’s dataset. (Y: available; N: not available)

**Table 3.** Summary of patients’ pathological scores for each cohort with mean and standard deviation. * denotes cohort-based pairwise significant differences (p value < 0.05).

|  | Lupus | ABMR | MR | TCMR |
| --- | --- | --- | --- | --- |
| Patients (n) | 23 | 9 | 15 | 9 |
| Tubulointerstitial Score (0-3) | 2.32 ± 0.80 | 1.40 ± 1.02 * | 2.53 ± 0.62 | 2.40 ± 0.66 |
| Chronicity Index (G + TI) (0-12) | 4.80 ± 3.26 | 3.60 ± 2.94 | 3.93 ± 1.61 | 4.50 ± 1.96 |
| Chronicity Index (G) (0-6) | 1.52 ± 1.78 | 1.00 ± 1.18 | 0.73 ± 0.85 | 0.70 ± 0.64 |
| Chronicity Index (TI) (0-6) | 3.28 ± 1.67 | 2.60 ± 2.01 | 3.20 ± 1.22 | 3.80 ± 1.89 |
| Tubule Fibrosis (TF) (0-3) | 1.64 ± 0.90 | 1.30 ± 1.00 | 1.60 ± 0.61 | 1.90 ± 0.94 |
| Tubule Atrophy (TA) (0-3) | 1.64 ± 0.81 | 1.30 ± 1.00 | 1.60 ± 0.61 | 1.90 ± 0.94 |

### Kidney structure segmentation and analysis pipeline

We first aimed to use machine learning to automatically identify different structural compartments of the kidney in a process known as instance segmentation. This task generated precise outlines, or “masks”, around individual objects of interest within a given image – an essential step for accurate, high-throughput medical imaging analysis. For this task, we trained three U-Net deep convolutional neural networks (Table 4) to segment instances of glomeruli, tubules, and capillaries (Fig. 1A). Training data consisted of whole-slide fluorescence raw images of canonical structural biomarkers (CD10, CD31, Claudin1 and DAPI for glomeruli; CD10, MUC1, Claudin1, and CD138 for tubules; CD31 for capillaries) with data processing steps reported in Methods. The training parameters were reported in Table 4 for each network, and training and testing datasets were generated by two human experts (Table 5). Instances for training and testing were selected randomly and distributed equally among normal and diseases cohorts. An Omnipose framework, which is a U-Net architecture designed for segmenting bacteria, was used to train for the segmentation tasks of kidney tubules and capillaries, as these structures are more asymmetrical and irregular in shape compared to the glomeruli (32, 39).

**Table 4.** Summary of the training parameters for the neural networks.

|  | Tubules | Glomeruli | Capillaries |
| --- | --- | --- | --- |
| Markers used as input | CD10, MUC1, Claudin1, CD138 | CD10, CD31, Claudin1, DAPI | CD31 (subtracting CD138) |
| Model framework | An Omnipose U-NET | A basic U-NET | An Omnipose U-NET |
| Pathology included in training dataset | LuN, ABMR, TCMR, MR | LuN, TCMR, MR, KC | LuN, ABMR, TCMR, MR |
| Epochs | 1000 | 100 | 1000 |
| Learning Rate | 0.01 | 0.0001 | 0.01 |
| Training tile size | 160x160x1 | 256x256x3 | 320x320x1 |

**Table 5.** Summary of the training and testing sets for the neural networks.

|  | **Tubules** | **Glomeruli** | **Capillaries** |
| --- | --- | --- | --- |
| Training | 8 WSIs | 78 WSIs | 8 WSIs |
|  | 420 tiles | 1124 tiles | 36 tiles |
| Testing | 6 WSIs | 56 WSIs | 7 WSIs |
|  | 7011 instances | 289 instances | 11442 instances |

Next, we classified the output masks into types and states based on the calculated fluorescence intensities of relevant biomarkers and shape features (Fig. 1B). Instances of tubules were classified into types (proximal or distal; based on expression of CD10 and MUC1) and states (stressed, inflamed, stressed and inflamed, atrophic; based on CD45, Claudin1, and each instance’s circularity feature). For capillaries, all instances were classified into states (inflamed, proliferating, or both; based on CD45 and Ki67 expression). Due to the sparse and random distribution of glomeruli per biopsy, we did not classify instances of glomeruli. The classified single instances of tubules and capillaries were used for downstream spatial analyses, in combination with pathological scores reported by a renal pathologist.

### Segmentation networks accurately identified glomeruli, tubules, and capillaries

When tested on independent, manually annotated validation sets, all three segmentation networks accurately identified glomeruli, tubules, and capillaries across all control and disease cohorts. Representative examples of prediction masks overlaid on fluorescence images demonstrated the pixel accuracy and sensitivity across different diseases and biopsy regions (Fig. 2). To quantify network performance, we computed F1 scores (on a scale from 0-1, where 1 indicates perfect precision and recall) (41). The average F1 score was 0.9 at an intersection over union (IoU) threshold of 0.5 (Fig. 2A-C, Table 5). All three networks performed best on the KC cohort, which had minimal regions of damage. While the F1 scores were lower in disease conditions compared to KCs, all networks maintained an F1 score above 0.8 at IoU threshold of 0.5. In particular, the glomeruli network had the lowest performance in LuN and ABMR cohorts, consistent with the widespread occurrence of glomerulonephritis in these diseases (55, 56) (Fig. 2A). Tubule and capillary networks showed negligible differences in performance across pathologies (Fig. 2B, 2C). In Supplementary Figure 1, additional performance metrics (precision, recall, and IoU mean as functions of IoU threshold) demonstrated comprehensive assessment of sensitivity and specificity of the three networks in detecting instances of glomeruli, tubules, and capillaries. In addition, when comparing network prediction masks and validated, ground-truth instances generated by human experts, the networks could distinguish closely situated structures in relatively dense tissue regions (Supplementary Fig. 1). These data reveal consistent and robust performance of the three networks. The occasions where they failed to predict accurately were usually at tissue edges where incomplete structures occur. Small regional mismatches between prediction and ground-truth also occured, which contributed to the decreasing performance at exceedingly high IoU threshold (> 0.8). However, in these cases, the minimal pixel mismatch did not impact the analysis at the object level. Overall, we identified a total of 1368 glomeruli, 911,352 tubules, and 443,260 capillaries across the disease cohorts in our high-dimensional immunofluorescence dataset (Table 6).

**Figure 2.**
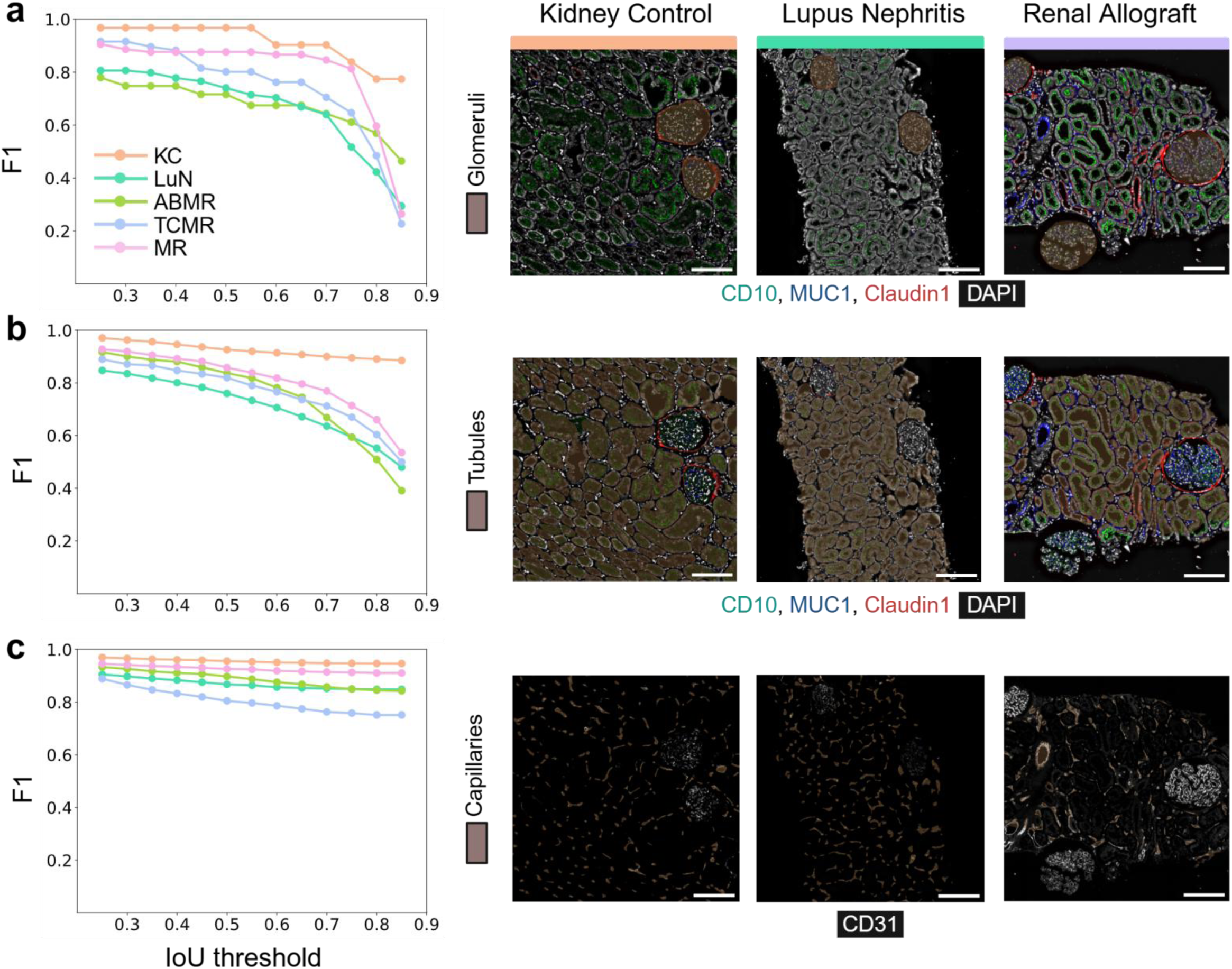
RDDx’s performance evaluated on an independent test set generated by human experts. F1 scores for instance segmentation performance across a range of intersection over union (IoU) threshold from 0.2 to 0.85 were plotted for all control and disease cohorts, along with a qualitative example for (a) glomeruli predictions, (b) tubules predictions, and (c) capillaries predictions.

**Table 6.**
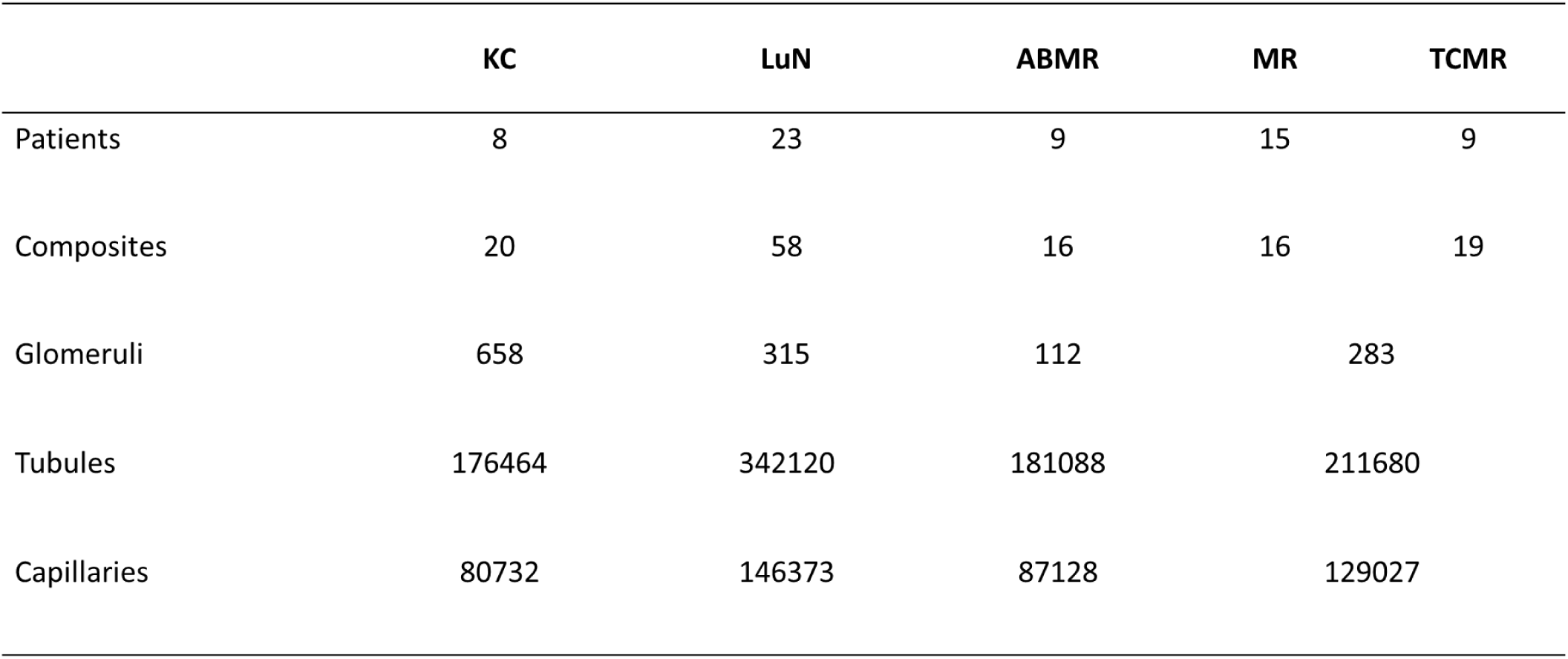
Summary of all structures identified and analyzed in this study.

When tested on a different in-house kidney CODEX dataset in which the CD31 marker for capillaries was replaced by CD34 (another endothelial marker), all three networks maintained robust performance without fine-tuning, as verified both quantitatively (average F1 score of 0.8 at 0.5 IoU) and qualitatively against human ground-truth (Supplementary Fig. 2). It is notable that the tubule and capillary networks resulted in more false negatives, as suggested by the drops in recall metrics and reduced accuracy of mask overlays, especially in less dense regions (Supplementary Fig. 2D, 2E). This suggests that human experts may fail to identify structures in regions with more damage, increased interstitial space, and fewer healthy structures.

### Quantifying structural features revealed different pathological states

Using segmentation masks from the three neural networks, we compared densities (areas) of different tubular and capillary states across the KC, LuN, and RAR cohorts. There were multiple statistically significant differential structural features across the cohorts (Fig. 3A). As expected, there were several differences between disease cohorts and controls. However, there were also striking differences between LuN and RAR, including differences in capillary states and elevated densities of stressed and inflamed tubules in the RAR cohort.

**Figure 3.**
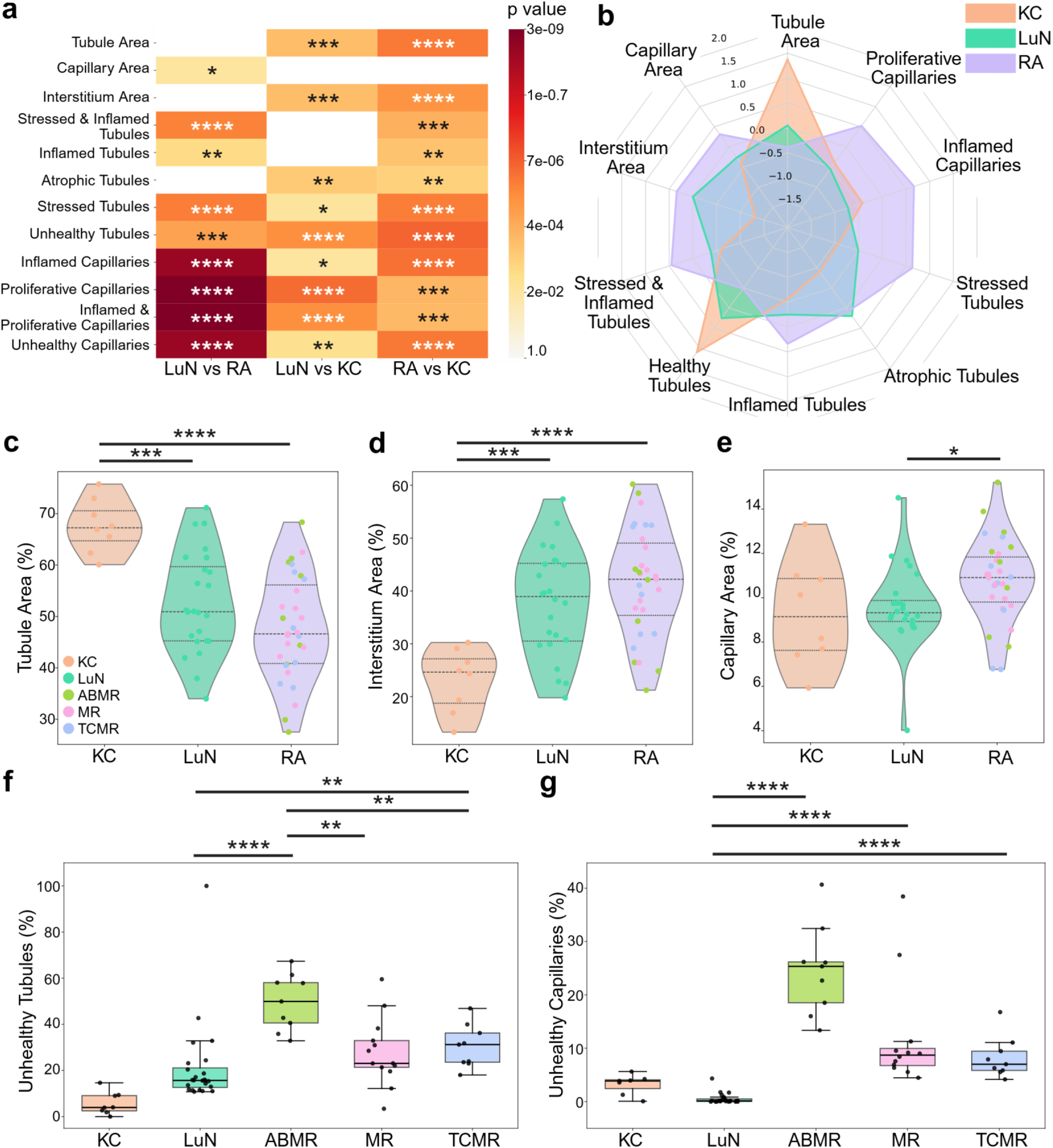
Biopsy-level structural compartment analysis characterizes differential structural changes and damages in lupus nephritis and renal allograft samples. (a) Statistical analysis (Mann Whitney U test for multiple two-sided pairwise comparisons with Benjamini-Hochberg corrections; α = 0.05) between unhealthy and control samples and between diseases. (b) A radar plot displays normalized compartment analysis of lupus nephritis, renal allograft, and kidney control. Violin plots show the distributions of each compartment by area for tubule (c), interstitium (d), and capillary (e). Each dot represents one biopsy. Boxplots comparing unhealthy tubule states (f) and unhealthy capillary states (g) across all cohorts: KC, LuN, ABMR, MR, and TCMR. All statistical analyses were done using Mann Whitney U test for multiple two-sided pairwise comparisons with Benjamini-Hochberg corrections; α = 0.05; p-values were reported as * for p < 0.05, ** for p < 0.01, *** for p < 0.001, **** for p < 0.0001.

To capture the distinct characteristics of each cohort, we overlaid three radar plots for KC, LuN, and RAR (Fig. 3B). In these plots, each radial line represents the indicated structural parameter scaled to have the same distribution range. As expected, the KC cohort showed the highest percentages of tubule area and healthy tubules and the lowest amount of interstitial space due to the tightly packed nature of tubules in a healthy kidney. Both LuN and RAR cohorts had elevated interstitial and atrophic tubule densities. Notably, the RAR cohort had significantly higher density of total capillaries, proliferating capillaries, and inflamed capillaries than LuN. This is consistent with the known role of microvascular injury in RAR (50, 56).

We next examined the distribution of different structural densities in individual biopsies. For these analyses, RAR was sub-categorized by biopsy into ABMR, MR, and TCMR (Fig. 3C-E). There was a progressive loss of tubular density from KC to LuN to RAR (Fig. 3C). Along with the loss of tubular density, there were increases in interstitium density in diseased cohorts (Fig. 3D). These relationships reflect that tubules and interstitium make up the majority of the tubulointerstitial space. Comparing the two disease cohorts, RAR had significantly more capillary densities than LuN (Fig. 3E). Interestingly, there appeared to be no correlation between RAR disease subsets and total capillary density. In addition to examining total densities by structure, we also examined densities of the unhealthy tubules and capillaries (Fig. 3F, 3G). The ABMR cohort had the highest density of both unhealthy tubules and capillaries. It is noted that the LuN cohort had a lower percentage of unhealthy tubules compared to the RAR subsets (Fig. 3F). Interestingly, the mean unhealthy capillary density in ABMR biopsies far exceeded MR and TCMR, with MR cohorts having two outliers that exhibited high unhealthy capillary densities (Fig. 3G). The observed MR heterogeneity is consistent with this disease category containing features of both ABMR and TCMR (14, 56).

### LuN biopsies had clustered regions of inflamed tubules

We next sought to understand the heterogeneous distributions and spatial patterns of healthy and diseased tubular states in our dataset. The KC cohort had the lowest percentage of unhealthy tubular states, followed by LuN, and then RAR (Fig. 4A). Mean densities of stressed, inflamed, and stressed and inflamed tubules consistently decreased across the three cohorts. This suggests that tubular stress is more prevalent than those leading to inflammation and immune cell invasion. As illustrated in a representative RAR biopsy (Fig. 4B), stressed tubules were the most prevalent and were often surrounded by CD45+ immune cells. However, CD45+ immune cells were present in a subset of tubules classified as inflamed. Interestingly, KC samples showed hardly any atrophic tubules, unlike the disease cohorts where atrophic tubules were scattered in regions containing many unhealthy tubules. These observations suggest that atrophy may be a more specific marker of disease.

**Figure 4.**
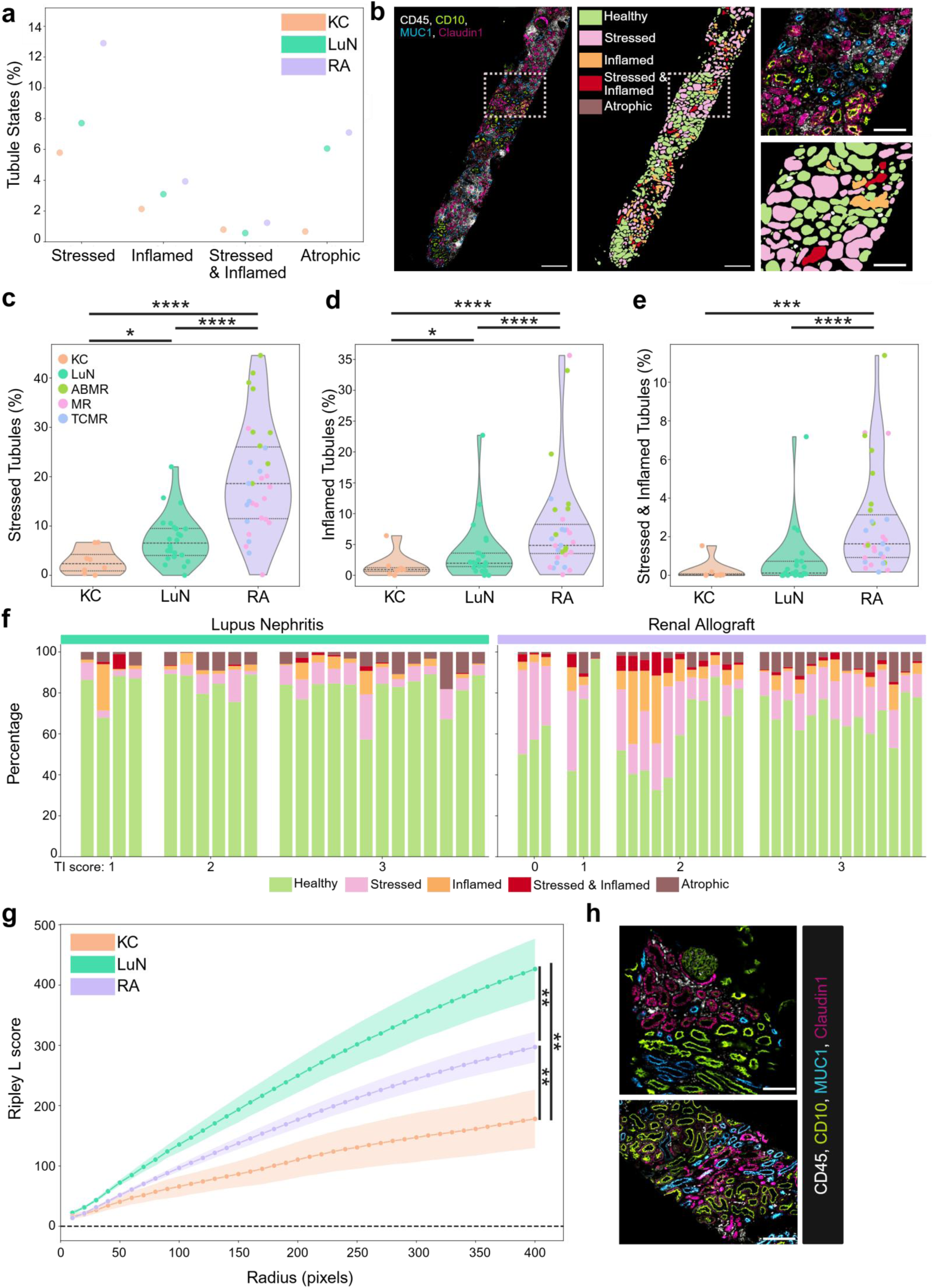
Tubular analysis highlights clustering patterns of unhealthy tubules in LuN. (a) Dot plots display the breakdown of tubular states for three datasets: KC, LuN, and RAR. (b) An example of a highly inflamed LuN sample with the fluorescence image and the tubule masks color-coded by state. Scale bar = 150 um. Violin plots show the distributions of stressed (c), inflamed (d), and stressed and inflamed tubules (e) across diseases and highlight the prevalence and significance of tubule states in the three RAR subsets. (f) Patient-level tubule state distributions (each patient was represented by a stacked bar) as grouped by pathologist-assigned tubulointerstitial inflammation score. (g) Spatial statistics analysis across the three datasets is calculated and plotted as Ripley L score across a range of radius from 0 to 400 pixels at a step of 10 pixels in the whole tissue biopsy. The shaded region denotes standard deviation when considering all tubular states per dataset. Two examples of spatially clustered inflamed and stressed tubules are shown in (h). Scale bar = 75 um. All statistical analyses were done using pair-wise Mann Whitney U tests for multiple two-sided comparisons with Benjamini-Hochberg corrections. p-values are reported as * for p < 0.05, ** for p < 0.01, *** for p < 0.001, **** for p < 0.0001.

Next, we analyzed the distributions of stressed, inflamed, and stressed and inflamed tubules across our cohorts (Fig. 4C, 4D, 4E). As expected, the KC cohort had very low percentages of unhealthy tubular states. Our classification was sensitive enough to detect a small subset of stressed tubules in KC biopsies suggesting regions surrounding renal cell carcinoma were not completely normal. Consistent with Fig. 3A and 3F, we observed significantly higher percentages of unhealthy tubules in RAR, compared to the LuN cohort. Furthermore, within the RAR cohort, ABMR biopsies tended to have higher densities of unhealthy tubules.

We next examined the distributions of tubules by state for each biopsy (Fig. 4F), organized by disease and pathological tubulointerstitial inflammation (TI) score (range 0-3) (12). In LuN, there were fewer unhealthy tubules overall compared to RAR. However, analysis of distributions of different unhealthy states by biopsy, LuN biopsies had a higher proportion of atrophic tubules, even in biopsies scored as mildly inflamed (TI score = 1). Interestingly, stressed tubules were frequently observed in biopsies with a TI score of 0 for RAR and were present at a lower percentage overall in LuN samples. RAR biopsies with the highest densities of inflamed and stressed or inflamed tubules had TI scores of 2. In RAR biopsies with a TI score of 3, indicating that in the cases with the most severe inflammation, the percentage of inflamed and stressed tubules declined while the proportion of atrophic tubules increased (Fig. 4F).

By our quantitative measures, LuN involved less inflammation in tubules (Fig 4d??). However, a similar number of LuN and RAR biopsies were scored severely inflamed (TI score of 3; Fig 4f). To explore this apparent discrepancy further, we calculated the spatial statistics (Ripley L) to determine the spatial patterns (clustering versus scattering) of unhealthy tubules across all datasets (Fig. 4G). The highest Ripley L score was observed in the LuN cohort, suggesting that unhealthy tubules were in patchy clusters. This pattern was evident in representative images of two LuN biopsies (Fig. 4H). In contrast, unhealthy tubules in the RAR cohort tend to cluster less, suggesting a more dispersed pattern of inflammation.

### ABMR biopsies manifested primary microvascular injury and capillary proliferation

Within the TI, most capillaries are peritubular. When analyzing this capillary compartment, ABMR biopsies had significant increases in capillary total density and the proportion of unhealthy capillaries (Fig. 3E, 3G). Therefore, we next compared the size of capillaries in KC and each RAR subset for both healthy and inflamed capillaries (Fig. 5A). Across cohorts, there was no significant difference in the size of healthy capillaries. In contrast, inflamed capillaries in both MR and TCMR were much larger. Interestingly, even though ABMR biopsies had far more inflamed capillaries (Fig. 3G), they were not as large as those in MR or TCMR. Inflamed capillaries (CD45+), proliferating capillaries (Ki67+), and double positive (CD45+Ki67+) capillaries were all elevated in the RAR cohort, particularly for the ABMR and MR subsets (Fig. 5B, 5C, 5D). It was notable that inflamed and proliferating capillary densities in LuN were reduced compared to KC and RAR cohorts and were not detectable in almost all biopsies. This indicates that, in our cohort, peritubular capillaritis is not a feature of LuN.

**Figure 5.**
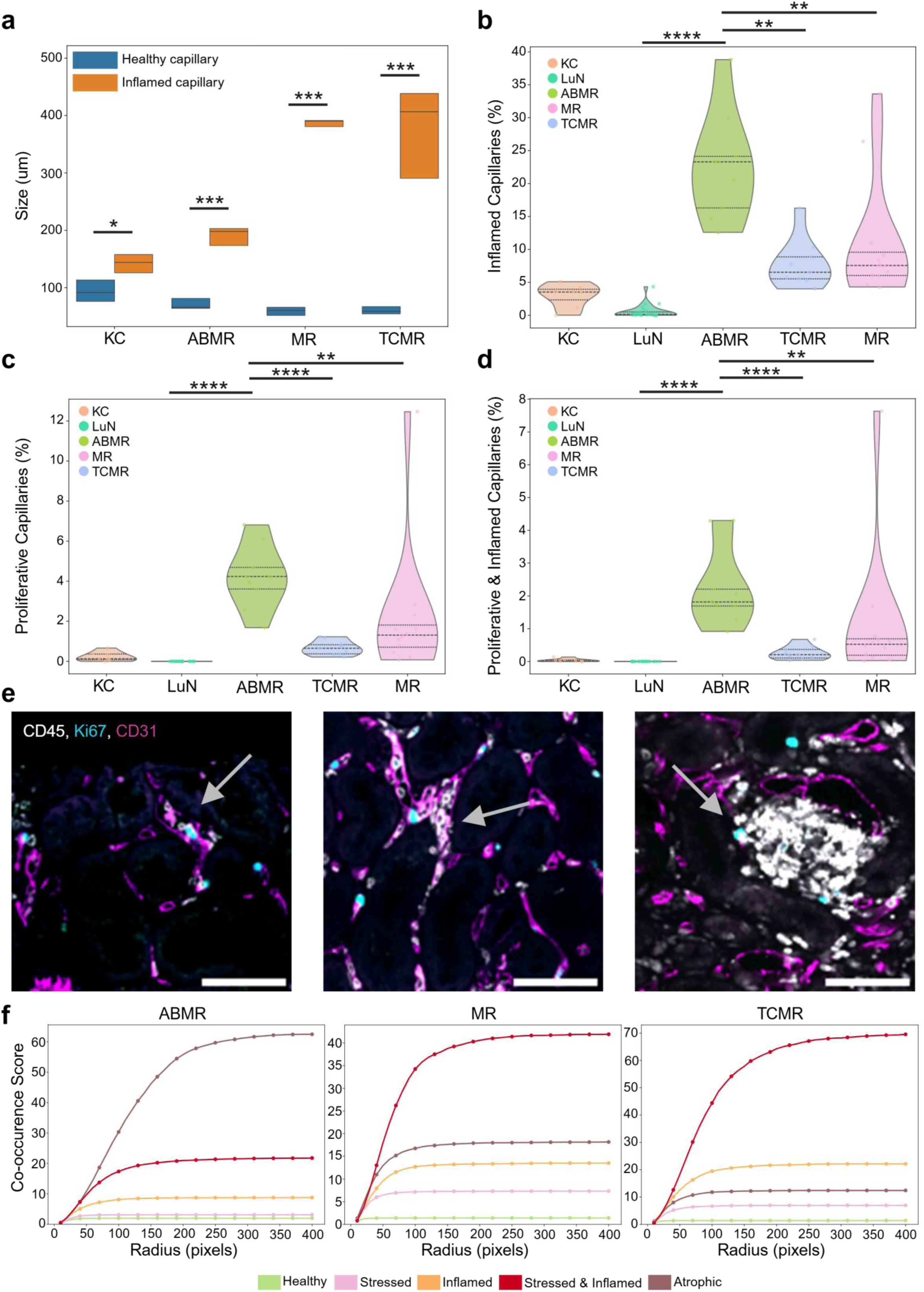
Capillary analysis highlights microvascular injury and increased capillary density in ABMR. (a) Size comparison between healthy and inflamed capillaries shown across kidney control and three RAR subsets. Violin plots show the distributions of inflamed capillaries (b), proliferating (c), and inflamed proliferating capillaries (d) across all control and disease cohorts. (e) Corresponding examples of inflamed capillaries and proliferating capillaries from each RAR subset are shown (CD31 in magenta, CD45 in white, Ki67 in cyan). Scale bar = 75 um. Arrows denote regions of unhealthy capillaries. (f) Co-occurrence scores between inflamed capillaries and each of tubule states in three RAR subsets are plotted for a range of radius from 0 to 400 pixels at a step of 10 pixels. All statistical analyses were pair-wise Mann Whitney U tests for multiple two-sided comparisons with Benjamini-Hochberg corrections. p-values are reported as * for p < 0.05, ** for p < 0.01, *** for p < 0.001, **** for p < 0.0001.

The finding of increased inflamed capillary density, but not size, in ABMR suggests that new blood vessels could form in response to immune cell injury. Indeed, the examples in Fig. 5E suggest that, in ABMR, there were a larger number of small capillaries and more proliferating cells (Ki67+) within those capillaries. In contrast, in TMCR, there were larger inflamed capillaries with Ki67+ cells in the interstitial space near capillaries.

We then asked where these inflamed capillaries were located with respect to unhealthy tubule niches by plotting their co-occurrence scores for the three RAR subsets (Fig. 5F). In ABMR, inflamed capillaries were most likely to be located near atrophic tubules, whereas in TCMR and MR, inflamed capillaries were most likely to be located near stressed and inflamed tubules.

### Quantitative structural analysis correlated with histology-based pathological scores

Given that histopathological scoring is the current standard for diagnosis and staging of LuN and RAR, we analyzed how tubule and capillary quantitative analyses correlated with manual scoring by a pathologist (Fig. 6A, 6B, 6C, 6D). All unhealthy tubule states positively correlated with increasing severity across all six reported scores, whereas healthy tubule percentage negatively correlated in all disease cohorts. Fig. 6B highlighted the significant correlations between stressed and unhealthy tubules with all the pathological scores of chronicity index (CIG, CITI, and CI) and tubular damage in TCMR. In MR, unhealthy tubules increased with chronicity index (Fig. 6C). Interestingly, the correlation between atrophic tubules, stressed tubules, and tubulointerstitial inflammation (TI) score for ABMR was poor, while inflamed tubules and inflamed capillaries had significant positive correlations to TI (Fig. 6D).

**Figure 6.**
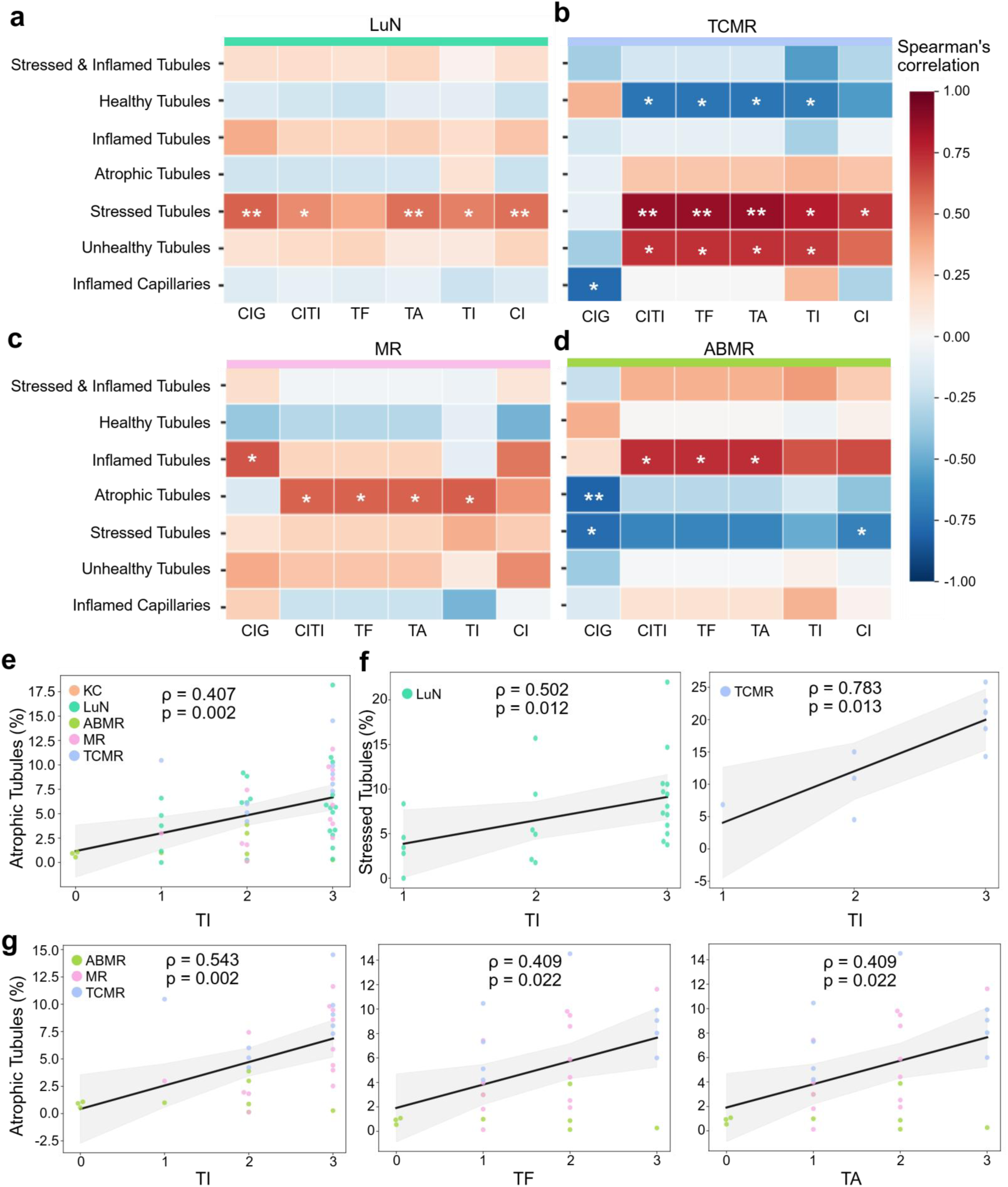
Structural analysis correlates with and complements pathological scores. Spearman’s correlations between quantitative structural analyses and categorical pathologist scoring of renal pathology: LuN (a), TCMR (b), MR (c), and ABMR (d). ρ coefficient values are displayed by color intensity while p values are reported as * for significant relationships. (e) Atrophic tubules percentage and tubulointerstitial inflammation (TI) score correlation across all datasets. (f) Stressed tubules percentage and TI score correlation in LuN and TCMR. (g) Atrophic tubules percentage and tubulointerstitial inflammation, interstitial fibrosis, and tubular atrophy correlations in three RAR subsets. All statistical analyses were pair-wise Mann Whitney U tests for multiple two-sided comparisons with Benjamini-Hochberg corrections. p-values are reported as * for p < 0.05, ** for p < 0.01, *** for p < 0.001, **** for p < 0.0001. CIG: chronicity index for glomerular inflammation (0–3); CITI: chronicity index for tubulointerstitial inflammation (0–3); TF: interstitial fibrosis (0–3); TA: tubular atrophy (0–3); TI: tubulointerstitial inflammation (0–3); CI: chronicity index (0–6).

Atrophic tubules were positively correlated to inflammation score across all datasets (p = 0.002) (Fig 6E). Using the inflammation score for specific disease cohorts, stressed tubules percentage significantly correlated with TI score in both LuN and TCMR (Fig. 6F). There were stressed tubules in biopsies with a TI score of 1 suggesting this is an early marker of injury. In the three RAR subsets, atrophic tubules were significantly correlated with tubulointerstitial inflammation, interstitial fibrosis, and tubular atrophy (Fig. 6G). These data suggest that tubular states can be a helpful quantitative complement to pathological scores.

## Discussion

Our study presents a novel, high-dimensional immunofluorescence dataset and computational pipeline (RDDx) to quantitatively assess kidney structures in healthy and disease states. To our knowledge these three trained deep convolutional neural networks are the first to achieve robust instance segmentation and classification of kidney structures and states based on high-dimensional whole-slide images. RDDx was highly accurate and demonstrated strong capacity to be generalized and applied to other datasets. Indeed, the sensitivity and specificity of RDDx is illustrated by the ability to accurately identify small, irregularly shaped structures, such as normal capillaries in densely infiltrated regions in LuN (16, 19). The precise identification of single instances of tubules and small capillaries allows us to differentiate tubulitis and capillaritis, both well-recognized predictors of adverse kidney outcomes (15,16). RDDx can be combined with cell phenotyping and molecular profiling to comprehensively map inflamed niches and *in situ* structural injuries.

Our study revealed distinct structural characteristics of LuN and RAR, confirming well-defined pathological features and identifying new ones for each disease. Both LuN and RAR showed decreasing tubule area with increasing interstitial space. While both LuN and RAR showed tubular structural damage and inflammation, RAR, especially ABMR, had significantly higher density of inflamed and proliferating capillaries compared to LuN, consistent with the known microvascular injury hallmark of ABMR (20, 38, 50). In contrast, LuN biopsies were marked by atrophic tubules, corresponding to reduced total tubular area and increased interstitial space. Additionally, LuN biopsies had clustered regions of inflamed tubules with the highest Ripley L scores, signifying spatially patchy tubular injury. This patchiness contrasted with the more diffuse injury pattern found in RAR, suggesting different injury patterns in autoimmunity and alloimmunity.

In LuN, the high prevalence of stressed tubules even in biopsies with low clinical inflammation scores, suggests that tubular stress might not be tightly linked to inflammation. Rather, the high prevalence of stressed tubules may be an adaptive response to inflammation (e.g. upregulation of tight junction proteins), consistent with previous observations of tubular epithelial cell stress signaling (45, 47). This stress response might be mediated by type 1 interferons which have been linked to progressive renal injury (23). We propose that chronic tubular stress, rather than direct invasion by inflammatory cells, leads to tubular atrophy and irreversible renal damage. Therefore, quantifying tubular atrophy might provide a precise, mechanistically informed, way to assess irreversible renal damage in LuN (45, 47, 48).

Within RAR subsets, ABMR featured distinct enrichment of inflamed capillaries and increased capillary density, whereas TCMR exhibited extensive tubular inflammation and stress. Notably, inflamed capillaries in ABMR were frequently found near atrophic tubules, suggesting that capillaritis was associated with hypoxic damage rather than tubular inflammation. In contrast, in TCMR capillaritis were in proximity to immune infiltrates and stressed tubules. This suggests that in TCMR, capillary inflammation was secondary to general tubulointerstitial inflammation. MR presented hybrid features of both—combining capillary inflammation of ABMR and tubular stress of TCMR—reflecting its mixed phenotype (50).

A key advance in our study was the quantification of proliferating capillaries, a feature not previously measurable in human histological renal sections. Traditional histology cannot capture proliferation markers like Ki67 within delicate capillary lumens, leading to this phenotype being overlooked. The spatial density of proliferating, inflamed, and double-positive (inflamed and proliferating) capillaries in ABMR suggests an important link between microvascular inflammation and allograft injury. This quantitative distinction between proliferating and inflamed capillaries provides new insight into potential neovascular and adaptive repair responses.

Our quantitative analyses demonstrated correlations between pathological tubular states (stressed, inflamed, atrophic) and conventional clinical histopathological scores, such as chronicity index, tubulointerstitial inflammation, fibrosis, and tubular atrophy. This suggests that RDDx measures similar features of disease activity and renal damage as clinical scoring. However, RDDx provides clear advantages over conventional scoring by a pathologist. For example, certain immune cell populations, particularly myeloid cells, are poorly visualized with conventional histological stains, leading to underestimation of inflammatory burden. In addition, manual scoring by pathologists can overlook heterogeneous or fibrotic regions and is subject to interobserver variability (14). Our quantitative analyses therefore complement conventional scoring by offering standardized, reproducible metrics.

Our study has limitations. Pipeline performance is subject to tissue quality and age. Archived biopsies that were collected over ten years ago tended to have reduced image quality, thus negatively impacting analysis. Although we were able to quantitatively analyze *in situ* tissue inflammation, we relied on highly curated, in-house CODEX datasets that are not routinely acquired in the clinic (44). In future studies, it will be important to develop a clinically useful tool by using high-dimensional data such as ours to develop techniques to identify and quantify different damage states in biopsies conventionally stained with H&E and periodic acid–Schiff (PAS).

Integrating our structural injury analyses with cellular phenotyping and molecular profiling will enable the delineation of immune cell states within renal regions and identify broad differences in the pathogenesis of different renal diseases. Furthermore, integration of immune states with precise assessments of structural damage will help identify different mechanisms of renal injury within each disease (45) and therefore parse mechanistic heterogeneity. Such investigations will also facilitate identifying biomarkers and designing novel therapeutic approaches (43).

In summary, our work establishes a quantitative and scalable deep learning–based pipeline (RDDx) to assess kidney damage in LuN and RAR. RDDx could be adaptable to other renal diseases such as diabetic nephropathy (54). Our studies reveal that each studied renal disease is associated with unique features of renal injury providing novel avenues for further mechanistic research and opportunities for developing better clinical tools for the diagnosis and staging of renal diseases.

## Figures

**Supplementary Figure 1. Additional instance segmentation performance metrics demonstrated accuracy of RDDx.** Precision, recall, and IoU mean across IoU threshold ranging from 0.2 to 0.85 were plotted for (a) glomeruli predictions, (b) tubules predictions, and (c) capillaries predictions. For each network, a whole-slide example of network prediction versus ground truth is shown, where consensus pixels are true positive, pixels as predicted by network only are false positive, and pixels as annotated by human experts only are false negative.

**Supplementary Figure 2. RDDx is generalizable and can predict on a different kidney CODEX dataset that uses CD34 as an endothelial marker (instead of CD31).** The zero-shot performance was evaluated: F1 score (a), IoU mean (b), precision (c), and recall (d). Fluorescence whole-slide image, along with network prediction versus ground truth overlays for each network, are shown in (e).

## Methods

### Data acquisition

All formalin-fixed paraffin-embedded (FFPE) core-needle kidney biopsies were sectioned at the thickness of 5 µm and imaged using the CODEX multiplexed imaging protocol described in (31). A panel containing eight biomarkers and DAPI for nuclei staining were used for this study (Table 1). Highly multiplexed immunofluorescence images were acquired of full biopsy sections on an Andor Dragonfly 200 Spinning Disk Confocal Microscope with the 40x objective lens and the pixel size of 0.1507 μm.

### Data preprocessing

Preprocessing steps of the highly multiplexed fluorescence images included stitching of tiles to create a full-section image and alignment of all fluorescence channels. Stitching and alignment were performed using Ashlar (40). Background subtraction and spectral normalization using min-max normalization to the 99th percentile was performed using tissue autofluorescence images collected at each wavelength used in data collection.

### Structural segmentation

We trained three U-Net based deep convolutional neural networks (DCNNs) to perform instance segmentation of glomeruli, tubules, and capillaries from whole-slide kidney fluorescence images. The U-Net architecture was selected due to its proven capability in biomedical image segmentation, specifically its encoder-decoder structure that captures multi-scale contextual information during the encoding phase. For the segmentation of tubules and capillaries—structures that exhibit higher morphological irregularity and asymmetry compared to glomeruli—an Omnipose variant of U-Net was implemented. Omnipose augments the U-Net backbone with flow field and distance field computations, enabling improved recognition of elongated and complex-shaped objects.

For the inputs to each segmentation network, a customized computational pipeline was applied to the normalized downsized fluorescence whole-slide images and canonical structural biomarkers of each structure were utilized to generate the input overlays: CD10, CD31, Claudin1, and DAPI for glomeruli; CD10, MUC1, Claudin1, and CD138 for tubules; and CD31 for capillaries. After the raw data collected from the microscope were stitched and aligned, these whole-slide images were normalized and pre-processed, followed by a downsizing step by a factor of 10.

Each DCNN was trained independently for their respective target structure, ensuring that feature extraction was optimally tuned to the morphology and biological features of that target. Model training parameters, including learning rate schedules, batch sizes, and optimizer settings, are specified in Table 4. The training and validation datasets were curated by two expert annotators (Table 5). Instance selections for training and testing were randomized and balanced to include equal representation from normal and diseased cohorts, mitigating potential bias from disease-specific biases.

### Structural classification

Instances of tubules are classified as proximal or distal based on 95th percentile marker expression patterns: Proximal tubules are identified by CD10 expression meeting or exceeding an intensity threshold of 20, while distal tubules are characterized by MUC1 expression at or above the same intensity threshold of 20. For tubules that remain unclassified after initial marker assessment, a secondary classification using the CD10:MUC1 expression ratio is applied. Tubules with a CD10:MUC1 ratio below 0.95 are reclassified as distal tubules, while those with ratios exceeding 1.05 are designated as proximal tubules. Tubular pathological states are determined as follows. Stressed tubules are identified when Claudin1 expression is above a 95th percentile pixel intensity of 45 with a frequency greater than 10. Inflamed tubules exhibit CD45 expression above 52 intensity with frequency exceeding 10. Tubules meeting criteria for both stressed and inflamed states are classified as "Stressed Inflamed." For tubules that are not classified for either proximal or distal types and do not fall into stressed, inflamed, or stressed inflamed categories, their circularity is calculated to determine whether they are atrophic tubules that downregulate both MUC1 and CD10 expressions. Atrophic tubules are characterized using morphometric parameters, specifically requiring size measurements above 69 pixels combined with circularity values at or above the 25th circularity percentile threshold (0.892805).

Instances of capillaries are classified as inflamed when their CD45 expression intensity of 50 or greater combined with a minimum CD45 frequency threshold of 5. To identify capillaries with proliferating cells, a Ki67 mask was generated for each biopsy using scipy.ndimage Gaussian blur and overlaid onto capillary mask. If there is an instance of Ki67 inside a capillary, that capillary is classified as proliferating. Inflamed proliferating capillaries are those that meet both criteria.

After obtaining all instance masks of kidney structures, we calculated the percentage of area occupied by tubules, vessels, and interstitium per biopsy by dividing the numbers of total pixels labeled as tubules, vessels, and interstitium over the total biopsy pixels. The size of each structure mask was calculated by transforming the total number of pixels labeled as that instance of interest into μm using the pixel size of 0.1507 μm.

### Spatial Statistics

To identify and quantitatively measure meaningful spatial patterns and behaviors of kidney structures, we performed two spatial statistics analyses: Ripley’s L and spatial cooccurrence.

To quantify the tendencies of clustering or dispersion of tubules at multiple spatial scales relative to complete spatial randomness for tubules in different data cohorts, we computed Ripley’s L score as a function of radius. First, spatial coordinates for centroids of each tubule were extracted from classified structural data, and pairwise Euclidean distances were calculated for each dataset-state group. For increasing radii from 0 to 400 pixels, the number of tubule pairs within each radius is counted and normalized by the sampling area, also known as the Ripley’s K statistic at each radius. Next, Ripley’s K values were transformed into Ripley’s L by L(r)=K(r)/π. The higher the Ripley’s L score, the higher level of clustering was observed in the dataset.

Spatial co-occurrence probability scores between different unhealthy tubule states and inflamed vessels were computed as a probability score: p(x∣y), indicating the probability of finding x given y. For every pair of x and y, we applied nearest-neighbor search to identify, for a range of radii from 0 to 400 pixels, the fraction of tubules of state y whose closest neighbor within the specified radius was an inflamed vessel x. This conditional probability score p(x∣y) was next normalized by the overall prevalence p(x) to get the co-occurrence score p(x∣y)/p(x), indicating the degree of spatial enrichment of tubules of state y around inflamed vessels.

### Statistical analysis

Statistical analysis was performed on Python. For all pair-wise comparisons, two-sided Mann-Whitney U tests were done with Benjamini-Hochberg corrections to decrease the false discovery rate. Significance as defined by p value was reported in the figure legends.

### Code availability

All scripts and models used for this study are available at https://github.com/ThaoCao/kidneystructureanalysis/.

All figures were assembled using Adobe Illustrator and BioRender.

### Data availability

Raw image data will be provided upon request.

## Acknowledgements

These studies were funded by the NIH Autoimmunity Centers of Excellence (AI082724), Department of Defense (LRI180083), Alliance for Lupus Research, Chan Zuckerberg Biohub, and NIH awards (S10-OD025081, S10-RR021039, and P30-CA14599). The authors are thankful for the University of Chicago Human Disease and Immune Discovery Core for facility, imaging, and computational support; Dr. Kevin Cutler for helpful suggestions on model training; Chun-Wai Chan for computational support; Drs. Jong Cheol Jeong and Rosario Perez Gutierrez for helpful insights on renal transplant rejection scoring criteria; Dr. Sarah May from Pipette & Quill Scientific Writing LLC for manuscript editing services, and the Gwen Knapp Center for Lupus and Immunology Research staff for additional support. All imaging acquisition was performed at the University of Chicago Human Disease and Immune Discovery Core. All raw data was subsequently processed on the MEL computational server in the Radiomics and Machine Learning Facility at the University of Chicago (256 Xeon Gold 6130 CPU cores, 3 TB of DDR4 ECC RAM memory, 24 TB of NVMe SSD storage, and 16 Nvidia Tesla V100 32GB GPU accelerators).

## Author Contributions

M.R.C. conceptualized the study. T.C., M.S.T., and G.C. conducted the data analysis. T.C. and J.A. acquired the data. S.H., J.A., and T.C. generated ground truth for machine learning model training. M.S.A. designed the data visualizer. M.L.G. provided computational resources. T.C. and M.R.C. wrote the manuscript. A.C. provided patient samples and clinical information. A.C. and A.C. contributed to results analysis. M.R.C. supervised the study. All the authors approved the final manuscript.

## References

1. StatPearls. End-Stage Renal Disease. StatPearls Publishing, Treasure Island, FL (2025).

2. De Pádua Netto, M. V. & Betônico, G. N. Lack of knowledge about chronic kidney disease and its consequences. Braz. J. Nephrol. 45, 134–135 (2023). doi:10.1590/2175-8239-jbn-2023-e004en

3. Bagavant, H. & Fu, S. M. Pathogenesis of kidney disease in systemic lupus erythematosus. Curr. Opin. Rheumatol. 21, 489–494 (2009). doi:10.1097/bor.0b013e32832efff1

4. Teoh, S. T., Yap, D. Y., Yung, S. & Chan, T. M. Lupus nephritis and chronic kidney disease: A scoping review. Nephrology 30, Article number: e14427 (2025). doi:10.1111/nep.14427

5. Chernova, I. Lupus nephritis: Immune cells and the kidney microenvironment. Kidney360 5, 1394–1401 (2024). doi:10.34067/kid.0000000000000531

6. Cornell, L. D., Smith, R. N. & Colvin, R. B. Kidney transplantation: Mechanisms of rejection and acceptance. Annu. Rev. Pathol. 3, Article number: 070912132245001 (2007). doi:10.1146/annurev.pathol.3.121806.151508

7. Cooper, J. E. Evaluation and treatment of acute rejection in kidney allografts. Clin. J. Am. Soc. Nephrol. 15, 430–438 (2020). doi:10.2215/cjn.11991019

8. Davidson, A. What is damaging the kidney in lupus nephritis? Nat. Rev. Rheumatol. 12, 143–153 (2015). doi:10.1038/nrrheum.2015.159

9. Palmer, S. C., Tumlin, J. A., Radhakrishnan, J., Rehaume, L., Cross, J., & Huizinga, T. The kidney injury biomarker profile of patients with lupus nephritis remains unchanged with the second-generation calcineurin inhibitor voclosporin. Front. Nephrol. 2, 1540471 (2025). doi:10.3389/fneph.2025.1540471.

10. Mas, V. R., Archer, K. J., Scian, M., & Maluf, D. G. (2010). Molecular pathways involved in loss of graft function in kidney transplant recipients. Expert Review of Molecular Diagnostics, 10(3), 269–284. 10.1586/erm.10.6

11. Sammaritano LR, Askanase A, Bermas BL, et al. 2024 American College of Rheumatology (ACR) Guideline for the Screening, Treatment, and Management of Lupus Nephritis. Arthritis Rheumatol. 2025;77(9):1115–1135. doi:10.1002/art.43212

12. Rojas-Rivera JE, García-Carro C, Ávila AI, et al. Diagnosis and treatment of lupus nephritis: a summary of the Consensus Document of the Spanish Group for the Study of Glomerular Diseases (GLOSEN). Clin Kidney J. 2023;16(9):1384–1402. Published 2023 Mar 22. doi:10.1093/ckj/sfad055

13. Fandel TM, Pfnür M, Schäfer SC, et al. Do we truly see what we think we see? The role of cognitive bias in pathological interpretation. J Pathol. 2008;216(2):193–200. doi:10.1002/path.2395

14. Schnuelle P. Renal Biopsy for Diagnosis in Kidney Disease: Indication, Technique, and Safety. J Clin Med. 2023;12(19):6424. Published 2023 Oct 9. doi:10.3390/jcm12196424

15. Londoño Jimenez A, Mowrey WB, Putterman C, Buyon J, Goilav B, Broder A. Brief Report: Tubulointerstitial Damage in Lupus Nephritis: A Comparison of the Factors Associated With Tubulointerstitial Inflammation and Renal Scarring. Arthritis Rheumatol. 2018;70(11):1801–1806. doi:10.1002/art.40575

16. Ding Y, Tan Y, Qu Z, Yu F. Renal microvascular lesions in lupus nephritis. Ren Fail. 2019;42(1):19–29. Published 2019 Dec 20. doi:10.1080/0886022X.2019.1702057

17. Clark MR, Trotter K, Chang A. The Pathogenesis and Therapeutic Implications of Tubulointerstitial Inflammation in Human Lupus Nephritis. Semin Nephrol. 2015;35(5):455–464. doi:10.1016/j.semnephrol.2015.08.007

18. Wilson PC, Kashgarian M, Moeckel G. Interstitial inflammation and interstitial fibrosis and tubular atrophy predict renal survival in lupus nephritis. Clin Kidney J. 2018;11(2):207–218. doi:10.1093/ckj/sfx093

19. Yaldır E, Cengiz BB, Açıkalın MF, Yaşar Bilge NŞ. The role of peritubular capillaritis in severity of lupus nephritis. Lupus. 2025;34(7):742–750. doi:10.1177/09612033251335821

20. Kozakowski N, Herkner H, Böhmig GA, et al. The diffuse extent of peritubular capillaritis in renal allograft rejection is an independent risk factor for graft loss. Kidney Int. 2015;88(2):332–340. doi:10.1038/ki.2015.64

21. Sultana, Z., Khatri, R., Yousefi, B. et al. Spatiotemporal interaction of immune and renal cells controls glomerular crescent formation in autoimmune kidney disease. Nat Immunol 26, 1977–1988 (2025). 10.1038/s41590-025-02291-8

22. Abraham R, Durkee MS, Ai J, et al. Specific in situ inflammatory states associate with progression to renal failure in lupus nephritis. J Clin Invest. 2022;132(13):e155350. doi:10.1172/JCI155350

23. Der E, Suryawanshi H, Morozov P, et al. Tubular cell and keratinocyte single-cell transcriptomics applied to lupus nephritis reveal type I IFN and fibrosis relevant pathways. Nat Immunol. 2019;20(7):915–927. doi:10.1038/s41590-019-0386-1

24. Samir V. Parikh, Ana Malvar, Huijuan Song, John Shapiro, Juan Manuel Mejia-Vilet, Isabelle Ayoub, Salem Almaani, Sethu Madhavan, Valeria Alberton, Celeste Besso, Bruno Lococo, Anjali Satoskar, Jianying Zhang, Lianbo Yu, Paolo Fadda, Michael Eadon, Dan Birmingham, Latha P. Ganesan, Wael Jarjour, Brad H. Rovin. Molecular profiling of kidney compartments from serial biopsies differentiate treatment responders from non-responders in lupus nephritis, Kidney International, Volume 102, Issue 4, 2022, Pages 845–865,

25. Rao, D.A., Arazi, A., Wofsy, D. et al. Design and application of single-cell RNA sequencing to study kidney immune cells in lupus nephritis. Nat Rev Nephrol 16, 238–250 (2020). 10.1038/s41581-019-0232-6

26. Arazi, A., Rao, D.A., Berthier, C.C. et al. The immune cell landscape in kidneys of patients with lupus nephritis. Nat Immunol 20, 902–914 (2019). 10.1038/s41590-019-0398-x

27. Casella G, Torcasso MS, et al. Immune cell quantification of in situ inflammation partitions human lupus nephritis into mechanistic subtypes. J Clin Invest. 2025;135(21):e192669. Published 2025 Sep 4. doi:10.1172/JCI192669

28. Keren L, Bosse M, Thompson S, et al. MIBI-TOF: A multiplexed imaging platform relates cellular phenotypes and tissue structure. Sci Adv. 2019;5(10):eaax5851. Published 2019 Oct 9. doi:10.1126/sciadv.aax5851

29. Lin JR, Izar B, Wang S, et al. Highly multiplexed immunofluorescence imaging of human tissues and tumors using t-CyCIF and conventional optical microscopes. Elife. 2018;7:e31657. Published 2018 Jul 11. doi:10.7554/eLife.31657

30. Goltsev Y, Samusik N, Kennedy-Darling J, et al. Deep Profiling of Mouse Splenic Architecture with CODEX Multiplexed Imaging. Cell. 2018;174(4):968–981.e15. doi:10.1016/j.cell.2018.07.010

31. Black, S., Phillips, D., Hickey, J. W., et al. CODEX multiplexed tissue imaging with DNA-conjugated antibodies. Nat. Protoc. 16, 3805–3835 (2021).

32. Ronneberger, O., Fischer, P., Brox, T. U-Net: Convolutional Networks for Biomedical Image Segmentation. Lecture Notes in Computer Science. 9351, 234–241 (2015). https://link.springer.com/chapter/10.1007/978-3-319-24574-4_28

33. Redmon, J., Divvala, S., Girshick, R., Farhadi, A. You Only Look Once: Unified, Real-Time Object Detection. Proceedings of the IEEE Conference on Computer Vision and Pattern Recognition. 2016, 779–788.

34. Stringer, C., Wang, T., Michaelos, M., & Pachitariu, M. Cellpose: a generalist algorithm for cellular segmentation. Nature Methods 18, 100–106 (2021).

35. Pachitariu, M., Stringer, C. Cellpose 2.0: how to train your own model. Nat Methods 19, 1634–1641 (2022). 10.1038/s41592-022-01663-4

36. Greenwald, N. F., et al. Whole-cell segmentation of tissue images with human-level performance using deep learning. Nature Biotechnology 40, 555–563 (2022).

37. Zappia, L. BANKSY: scalable cell typing and domain segmentation for spatial omics. Nature Reviews Genetics 25, 527–528 (2024).

38. Djamali, A., Kaufman, D. B., Ellis, T. M., Zhong, W., Matas, A., & Samaniego, M. Diagnosis and Management of Antibody-Mediated Rejection: Current Status and Novel Approaches. Am J Transplant. 14, 255–271 (2014).

39. Cutler, K. J., Stringer, C., Wiggins, P. A., & Hasty, J. Omnipose: a high-precision, morphology-independent solution for bacterial cell segmentation. Nature Methods 20, 292–299 (2023).

40. Muhlich, J. L., Chen, Y.-A., Yapp, C., Russell, D., Santagata, S., & Sorger, P. K. Stitching and registering highly multiplexed whole-slide images of tissues and tumors using ASHLAR. Bioinformatics 38, 4613–4621 (2022).

41. Reinke, A., Tizabi, M.D., Baumgartner, M. et al. Understanding metric-related pitfalls in image analysis validation. Nat Methods 21, 182–194 (2024). 10.1038/s41592-023-02150-0

42. Maragall, J., Lucarelli, N., Naglah, A., Border, S., Winfree, S., Laszik, Z., Eadon, M. T., El-Achkar, T. M., Jain, S., & Sarder, P. CODEX and H&E imaging: cell type mapping, analysis, and visualization pipeline. SPIE Medical Imaging, 31. (2024) 10.1117/12.3008471

43. Huang, Z., Yang, E., Shen, J. et al. A pathologist–AI collaboration framework for enhancing diagnostic accuracies and efficiencies. Nat. Biomed. Eng 9, 455–470 (2025). 10.1038/s41551-024-01223-5

44. Dayao, M. T., Mayer, A. T., Trevino, A. E., & Bar-Joseph, Z. (2025). Using spatial proteomics to enhance cell type assignments in histology images. Cell Reports Methods, 101204. 10.1016/j.crmeth.2025.101204

45. Abedini, A., Levinsohn, J., Klötzer, K. A., Dumoulin, B., Ma, Z., Frederick, J., Dhillon, P., Balzer, M. S., Shrestha, R., Liu, H., Vitale, S., Devalaraja-Narashimha, K., Grandi, P., Bhattacharyya, T., Hu, E., Pullen, S. S., Boustany-Kari, C. M., Guarnieri, P., Karihaloo, A.,… Susztak, K. (2022). Spatially resolved human kidney multi-omics single cell atlas highlights the key role of the fibrotic microenvironment in kidney disease progression. bioRxiv. 10.1101/2022.10.24.513598

46. Wu, H., Dixon, E.E., Xuanyuan, Q. et al. High resolution spatial profiling of kidney injury and repair using RNA hybridization-based in situ sequencing. Nat Commun 15, 1396 (2024). 10.1038/s41467-024-45752-8

47. Lake, B.B., Menon, R., Winfree, S. et al. An atlas of healthy and injured cell states and niches in the human kidney. Nature 619, 585–594 (2023). 10.1038/s41586-023-05769-3

48. Polonsky, M., Gerhardt, L.M.S., Yun, J. et al. Spatial transcriptomics defines injury specific microenvironments and cellular interactions in kidney regeneration and disease. Nat Commun 15, 7010 (2024). 10.1038/s41467-024-51186-z

49. Xuanyuan, Q., Wu, H., Sundaramoorthi, H. et al. Multimodal spatial transcriptomic characterization of mouse kidney injury and repair. Nat Commun 16, 7567 (2025). 10.1038/s41467-025-62599-9

50. Gibson, I.W., et al. Peritubular Capillaritis in Renal Allografts: Prevalence, Scoring System, Reproducibility and Clinicopathological Correlates. American Journal of Transplantation, Volume 8, Issue 4, 819–825

51. Garreta, E., Moya-Rull, D., Centeno, A. et al. Systematic production of human kidney organoids for transplantation in porcine kidneys during ex vivo machine perfusion. Nat. Biomed. Eng (2025). 10.1038/s41551-025-01542-1

52. Khoshdel-Rad N, Ahmadi A, Moghadasali R. Kidney organoids: current knowledge and future directions. Cell Tissue Res. 2022;387(2):207–224. doi:10.1007/s00441-021-03565-x

53. Liang J, Liu Y. Animal Models of Kidney Disease: Challenges and Perspectives. Kidney360. 2023;4(10):1479–1493. doi:10.34067/KID.0000000000000227

54. Liu DD, Hu HY, Li FF, et al. Spatial transcriptomics meets diabetic kidney disease: Illuminating the path to precision medicine. World J Diabetes. 2025;16(9):107663. doi:10.4239/wjd.v16.i9.107663

55. Najafi CC, Korbet SM, Lewis EJ, et al. Significance of histologic patterns of glomerular injury upon long-term prognosis in severe lupus glomerulonephritis. Kidney Int. 2001;59(6):2156–2163. doi:10.1046/j.1523-1755.2001.00730.x

56. Halloran PF, Merino Lopez M, Barreto Pereira A. Identifying Subphenotypes of Antibody-Mediated Rejection in Kidney Transplants. Am J Transplant. 2016;16(3):908–920. doi:10.1111/ajt.13551

