## Supplementary figures and images for "High-dimensional spatial proteomics and novel machine learning pipeline identifies disease specific renal damage states"

### Supplemental Figures

Cao, et al. Supplementary Fig 1

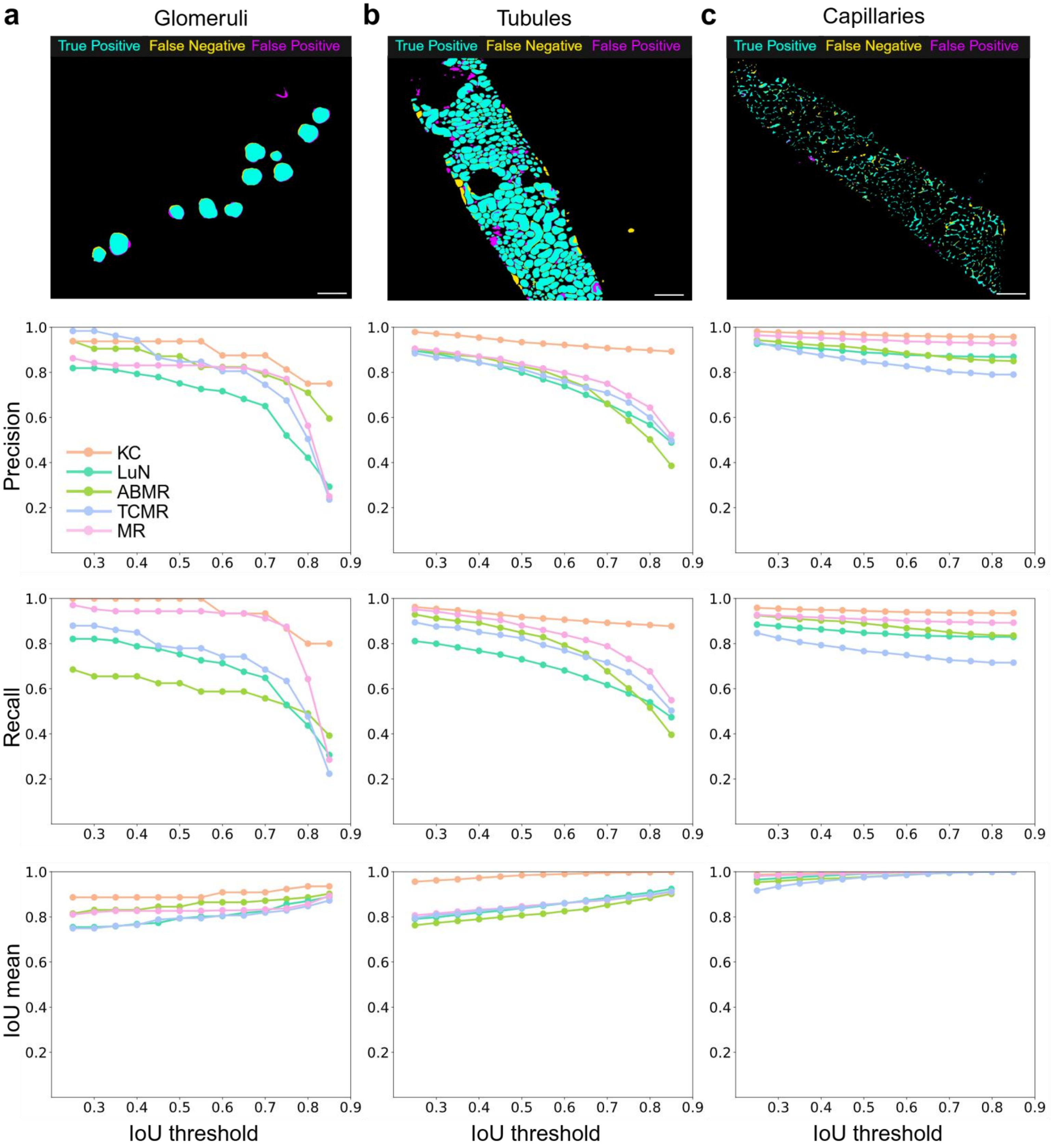

Cao, et al. Supplementary Fig 2

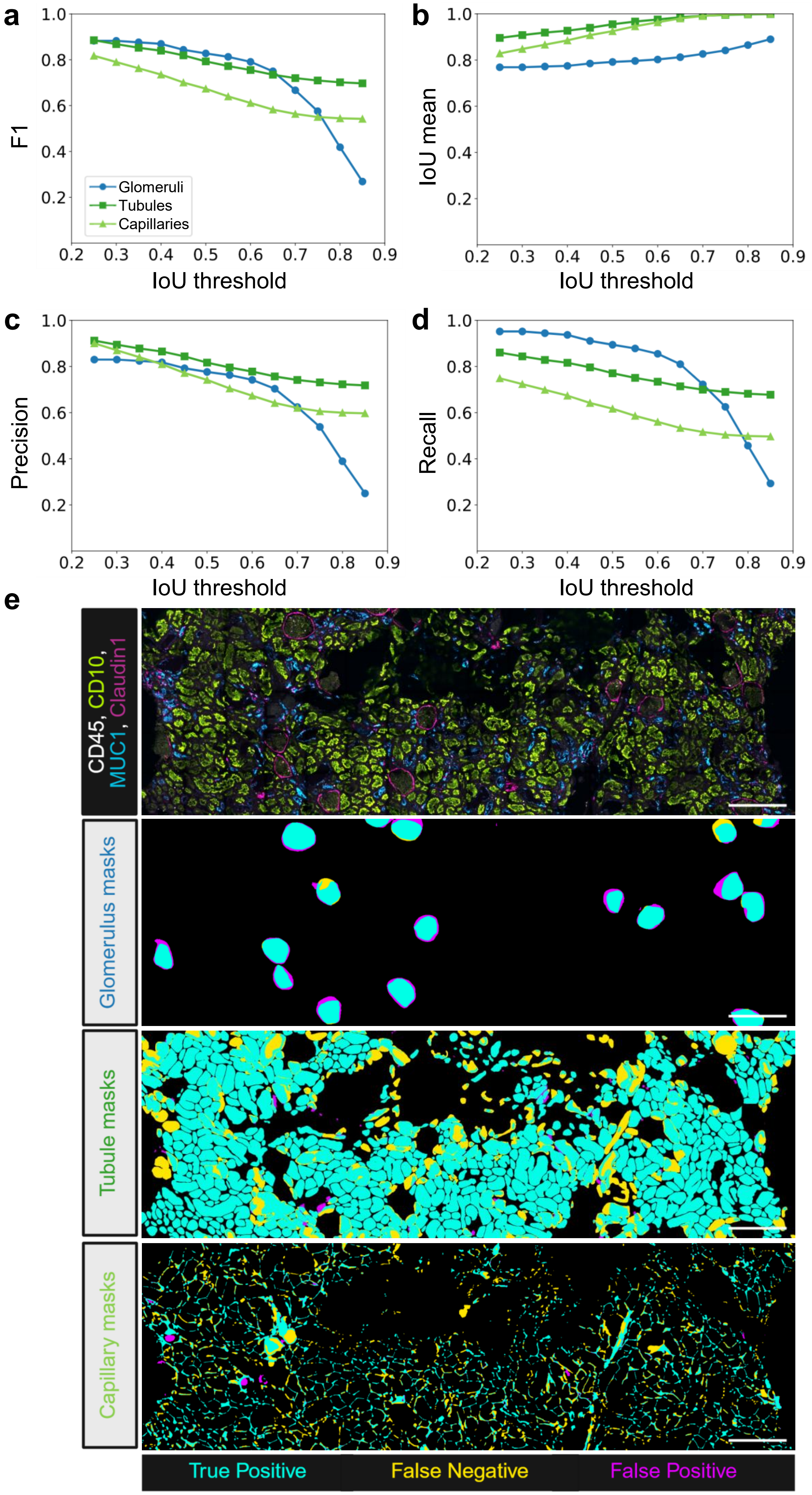
